# Modelling the dynamics of vesicle reshaping and scission under osmotic shocks

**DOI:** 10.1101/2020.11.16.384602

**Authors:** Christian Vanhille-Campos, Anđela Šarić

## Abstract

We study the effects of osmotic shocks on lipid vesicles via coarse-grained molecular dynamics simulations by explicitly considering the solute in the system. We find that depending on their nature (hypo- or hypertonic) such shocks can lead to bursting events or engulfing of external material into inner compartments, among other morphology transformations. We characterize the dynamics of these processes and observe a separation of time scales between the osmotic shock absorption and the shape relaxation. Our work consequently provides an insight into the dynamics of compartmentalization in vesicular systems as a result of osmotic shocks, which can be of interest in the context of early proto-cell development and proto-cell compartmentalisation.

## I. INTRODUCTION

Lipid membranes are one of the most important building blocks in cellular biology, forming the envelope of the cell and defining the structure and function of intracellular units in eukaryotic cells. Given their physio-chemical properties, biological membranes present very particular reshaping capabilities that are intimately related to the proper functioning of the cellular units they constitute. Cell membrane morphology transformations in nature can occur in a number of different ways, which can also include creating of new units such as budding or division. Budding is the process by which a portion of a membrane detaches, usually in the form of a vesicle that is smaller than the mother membrane. Such processes are ubiquitous in cellular biology, playing important roles in vesicle secretion, protein transport, organelle synthesis and endocytosis [1]. In the division, a lipid vesicle or a cell splits into two closed units of comparable sizes. In more evolved cells this process is usually very specifically driven by a set of proteins (such as actomyosin [2], ESCRT-III [3, 4] or FtsZ [5]) composing the divisome machinery, responsible for exerting the required forces on the membrane. However, division of vesicles can also occur in spontaneous ways usually caused by environmental changes (shear stress [6], osmotic shocks [7–9]), compositional changes to the membrane (protein addition [10, 11]) or even motility of nascent cells [12].

In this context, vesicles are especially relevant as they represent a typical model system to study membrane reshaping events. Moreover, in the context of early life evolution and proto-cells development in particular, vesicles are thought to have played a crucial role, having evolved towards the existing forms of cellular life by compartmentalizing and developing more complex behaviors as a result [13–15]. There exist a number of experimental, theoretical and numerical works that examine the connection of distinct vesicle reshaping events to their physio-chemical conditions. Previous experimental efforts have focused on understanding the role of the physical properties (e.g. fluidity [16], multiphase domain coexistence [17, 18]) and composition (e.g. protein adsorption at high [10, 19] and low [11] surface density) of membranes in reshaping Giant Unilamellar Vesicles (GUVs). Other experimental approaches have focused on the effect of osmotic shocks and other external forces (e.g. shear stress [6]) on vesicle morphology. It is thus known that hypertonic shocks (when the solute concentration is larger outside than inside the vesicle) induce strong deformations on GUVs with varying results depending on the specific composition and chemical properties of the membrane [9, 20, 21]. Division can thus occur when osmotic stress is coupled with phase separation in the lipid bilayer [7, 8]. More important for cell compartmentalization and endocytosis, simple osmotic shocks can also drive inward budding of the membrane coupled with absorption of external medium into the newly created inner compartment [9, 14, 15, 22, 23]. Finally, studies have also shown that hypotonic shocks result in pulsatile dynamics of stretch and release for the vesicle until equilibrium is reached between the two solutions [24, 25].

Regarding theoretical approaches, most advances came at the end of the 20th century, with the introduction of the Helfrich model of bending free energy [26]. Subsequently, most experimentally observed vesicles conformations were described by applying free energy minimisation principles focusing primarily on the membrane curvature, the so-called curvature models [27, 28]. Although very effective in describing equilibrium shapes, such theoretical approaches present some limitations. Probably the most important limitation comes from working with continuous surfaces, which makes it impossible to describe budding or membrane scission events, resulting in unrealistic limit shapes. Furthermore, given that usually such models assume constant surface area, no stretching of the membrane is allowed, rendering any hypotonic shock scenario virtually inaccessible. Indeed, hypotonic shocks result in a swelling and bursting of the vesicle sometimes displaying cyclic relaxation dynamics [24, 25] which cannot be replicated by a non-stretching surface description of the lipid bilayer that cannot break. Finally, by the very nature of the free energy minimisation approach, curvature models provide little insight into the relaxation dynamics of the vesicles.

An alternative approach are physiochemical kinetic models [29–31], which do offer a solution for vesicle volume dynamics, for instance, but do not capture the shape transformations associated with it. Lastly, existing numerical efforts in trying to describe vesicle shape transformations are usually limited to Monte Carlo simulations of meshed surfaces governed by energy considerations associated with curvature [32]. These are again successful in predicting most equilibrium shapes, but provide little help in overcoming the limitations presented by the theoretical models à la Helfrich. One exception to these examples, which manages to reproduce budding events, is the approach taken by Yuan *et al*. [33]. The authors used molecular dynamics simulations to study the effects of volume changes resulting from osmotic shocks on various types of vesicles. One limitation of this study is that, by explicitly controlling the volume dynamics via a prescribed protocol, an artificial relaxation rate was introduced [29, 30].

In the present work we focus on vesicle reshaping as a result of explicit osmotic shocks. In order to overcome limitations posed by the previous methods, where reshaping dynamics and vesicle breakage were inaccessible, we resort to using molecular dynamics simulations of coarse-grained model lipid bilayers under an explicit gradient of solute particles. This approach allows us to characterize the dynamic process leading to the different morphological transformations of vesicles. We find that hypertonic osmotic shocks can result in the engulfment of external material into inner vesicular compartments while hypotonic shocks on the contrary cause vesicle dilation and oscillatory bursting for sufficiently strong concentration differentials. We observe a separation of time scales in the dynamics of the shock absorption and shape transformations, where the vesicle first absorbs shocks by forming many local deformations and transient shapes, before relaxing to an equilibrium shape. Our study provides new insights into the dynamics leading to vesicle deformations and, in particular, vesicle endocytic events as a result of environmental osmotic stresses.

## II. SIMULATION DETAILS

In order to study the effect of osmotic shocks on lipid vesicles we carry out coarse-grained (CG) molecular dynamics (MD) simulations where we explicitly consider the solute modeled as CG particles. The system thus consists of a spherical model membrane surrounded by solute particles on the outside at a concentration *ρ*_*out*_ (see purple particles in Fig.1) and encapsulating inner solute particles at a different concentration *ρ*_*in*_ (see cyan particles in Fig.1). All particles are volume-excluded spheres of a diameter *σ, σ* being the MD unit of length, and mass *m* = 1 embedded in cubic box of constant volume *V* = *L*^3^ with *L* = 44 *σ* and within periodic boundary conditions. See Figure 1 for the snapshots of the system.

**FIG. 1:**
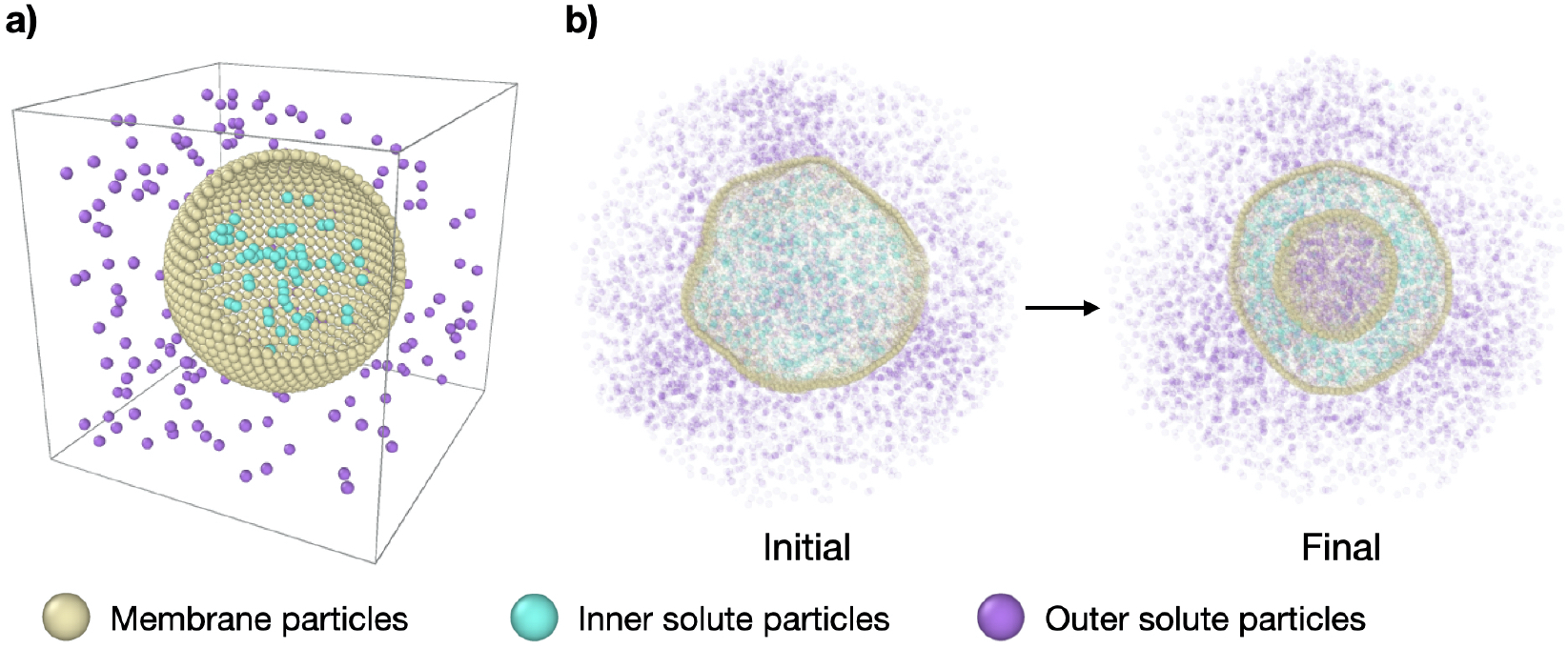
Molecular dynamics simulation setup. (a) Simplified representation of the system: a spherical vesicle of CG lipids (yellow particles) in a cubic box with solute particles inside (cyan) at a concentration *ρ*_*in*_ and solute particles outside (purple) at a concentration *ρ*_*out*_. (b) Cross-section snapshots of the simulated system at initial conditions (*τ* = 0 time steps) and equilibrium (*τ* = 3 × 10^5^ time steps) for representative parameters. The outer solute concentration is *ρ*_*out*_ = 0.18 and the concentration ratio is *ν* = 0.31. See Supplementary Movie 2b for an illustration of the transformation sketched in the right panel.

The membrane is modelled using the single particle thick model developed by Yuan *et al*. [34] which is capable of fission and fusion events and reproduces biologically relevant mechanical properties of membranes. In this model, bilayer sections of around 10 nm in width are represented by single beads interacting via an attractive potential. The bending energy is implemented via interactions between vectors on each bead, which represent the local normal of the membrane and are directly related to its curvature. The specific parameters of the model are chosen to encode for a fluid membrane of vanishing spontaneous curvature and bending rigidity of 15 *k*_*B*_*T, k*_*B*_ being the Boltzmann’s constant and *T* being temperature. For more details on the interaction potential used in this model and the particular values used here refer to Section VI in the Supplementary Information^†^. All other interactions are modeled as simple volume exclusion repulsive interactions using a Weeks-Chandler-Anderson potential.

The system is always initialised in the same way: i) *N*_*mem*_ = 4322 membrane particles define a relaxed spherical vesicle of mean radius *R* ∼ 17 *σ* at the center of the box and ii) solute particles take random positions on a hexagonal lattice at set concentrations *ρ*_*in*_ and *ρ*_*out*_ inside and outside of the vesicle, respectively, ensuring no overlap occurs with the membrane. In this way, the osmotic conditions of the system are fully defined by two parameters only: the outer solute concentration, *ρ*_*out*_, and the ratio of concentrations *ν* = *ρ*_*in*_*/ρ*_*out*_. The latter quantity is often referred to as the theoretical reduced volume. Hence, setting *ν <* 1 defines hypertonic conditions while *ν >* 1 corresponds to hypotonic conditions. In this setup the vesicles studied have a diameter of a few hundred nanometers, which is within the range of physiologically relevant scales. However, to test the role of the vesicle size, we also explored larger model vesicles (see Section VII in the Supplementary Information^†^). We run simulations using the open source molecular dynamics package LAMMPS [35] with a Langevin thermostat (at reduced temperature *T* = 1 and damping coefficient of 1) within the *NV E* ensemble, *E* being the total energy of the system, for 3 × 10^5^ time steps, each of size 0.01 MD time units (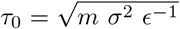 with *ϵ* = *k*_*B*_*T* the MD energy unit).

## III. RESULTS

### A. Characterising osmotic shocks

First we make sure that explicitly simulating solute particles in a closed system with a coarse-grained model lipid vesicle properly reproduces osmotic shocks. For this we measure the initial osmotic pressure measured across the vesicle (see Section I in the Supplementary Information^†^ for details on this). Figure 2a presents the dependence of the osmotic shock on the two system parameters: the outer particle absolute concentration, *ρ*_*out*_, and the ratio of concentrations between the vesicle interior and exterior, *ν* = *ρ*_*in*_*/ρ*_*out*_. Figure 2b shows the fit agreement between the measured osmotic pressure with the expected value of the osmotic pressure, as given by the equation Π = *RT ρ*_*out*_(*ν* − 1) [27]. Indeed, beyond a transformation factor due to MD units, we find that we are truly applying an osmotic shock to the vesicle and its magnitude scales as expected theoretically.

**FIG. 2:**
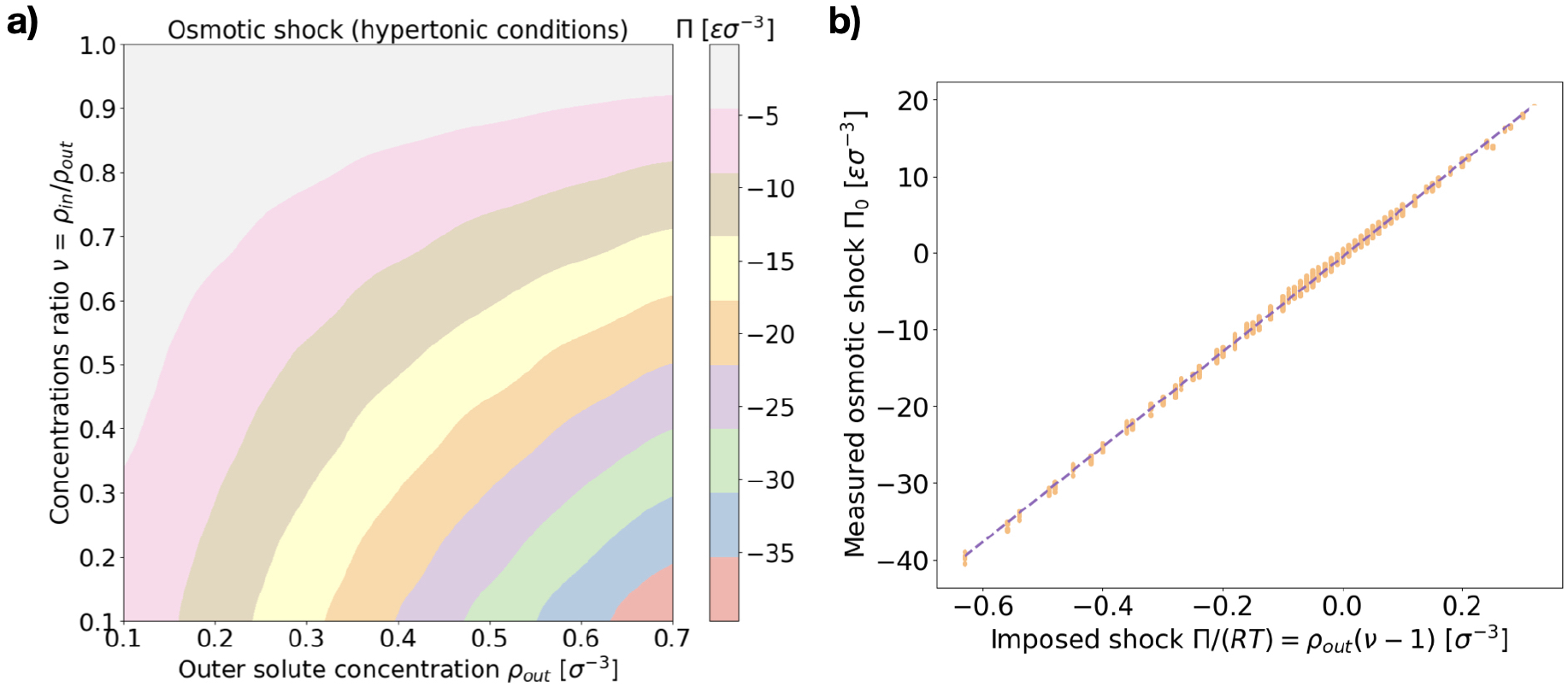
Osmotic shock at initial conditions. (a) Measured values of the initial osmotic pressure difference across the membrane (osmotic shock) Π (color coded) as a function of the two system parameters {*ρ*_*out*_, *ν*} for hypertonic conditions *ν <* 1. Values are in MD units of pressure: [Π] = *ϵσ*^−3^. (b) Measured values of the osmotic shock (orange dots, again in MD units) against the corresponding expected theoretical value Π*/*(*RT*) = *ρ*_*out*_(*ν* − 1). A linear fit is included as well (dashed line). The left panel shows the average over 10 independent runs for each parameter set while the right panel shows all the individual measurements.

### B. Coarse-grained model membranes recover known equilibrium shapes

We are next interested in investigating whether our system is able to reproduce the relaxation transformations observed experimentally on GUVs [9, 15], predicted theoretically [26, 28], or simulated by other means [32, 33]. We thus look at the equilibrium shapes our simulated vesicles present in hypertonic conditions (larger solute concentration outside than inside the vesicle). While the spectrum of shapes and different morphologies lipid vesicles present when subject to hypertonic shocks is vast and not necessarily restricted to a few discrete options, we can nonetheless categorise our results into several distinct kinds of shapes that help analyze the results. Inspired by previous works using similar classifications [15, 28, 32] we distinguish six different equilibrium vesicle morphologies according to which we classify our results (see Figure 3 and accompanying snapshots):

**FIG. 3:**
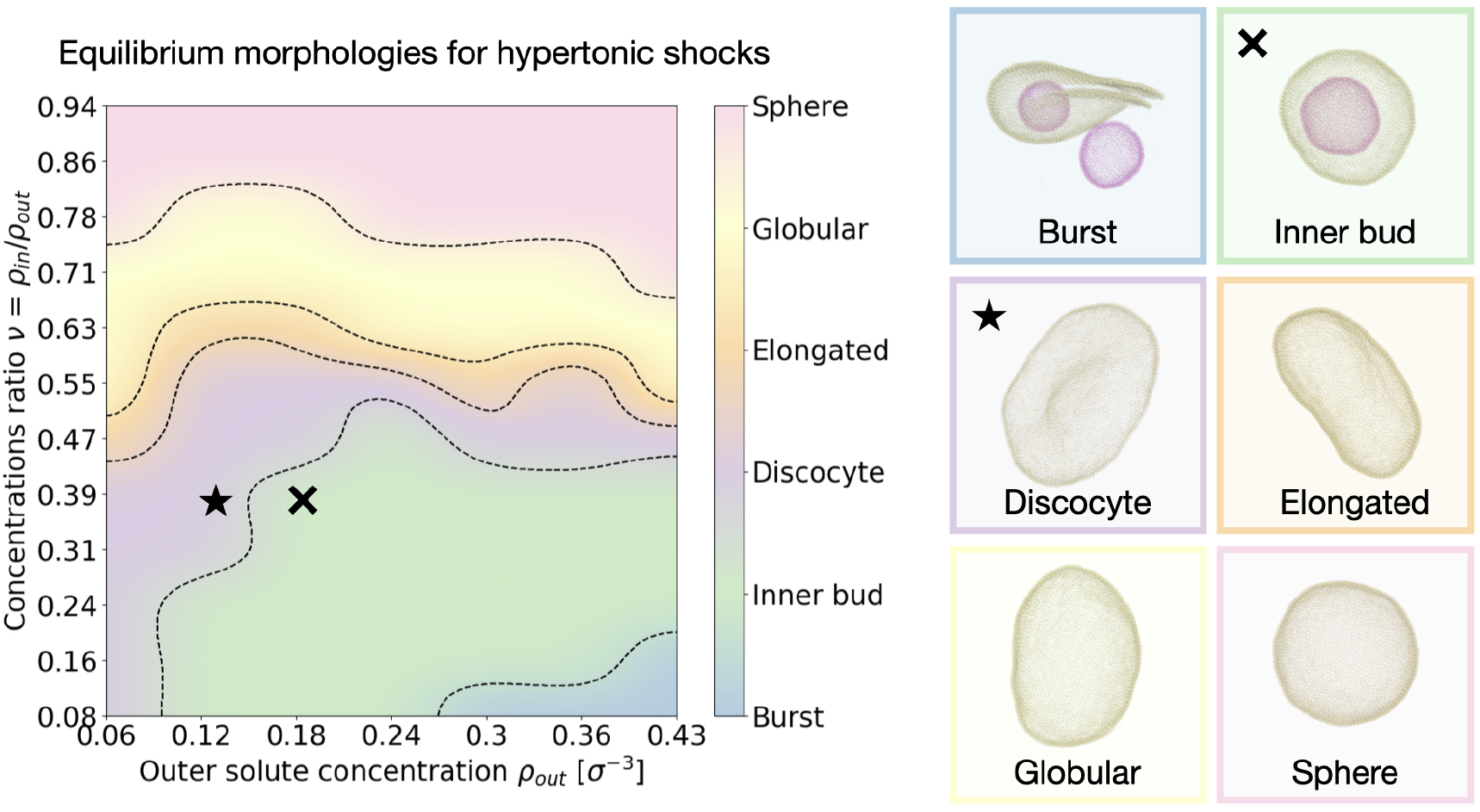
Final morphology diagram for hypertonic conditions. We distinguish six different characteristic equilibrium shapes of the vesicle, represented to the right of the figure. The resulting observed morphology for each point in the parameter space {*ρ*_*out*_, *ν*} is color coded accordingly. Dotted lines indicate transitions. The cross and star symbols refer to the dynamics curves below (see Figures 4 and 5). Snapshots are framed using the same color code as the diagram on the left. Particle coloring is used to distinguish multiple vesicles in the same system. All morphologies correspond to equilibrium, reached after *τ* = 3 *×* 10^5^ time steps.

1. *Burst*: at high outer solute concentration *ρ*_*out*_ and very low ratio of inner and outer concentrations *ν*, the osmotic shock causes the mother vesicle to burst, often yielding several smaller vesicles containing a mixture of both inner and outer solutions (Supplementary Movie 2a).
2. *Inner bud*: for a large subset of the parameter space where the inner to outer concentration ratio is still small (*ν <* 0.5), but the shock is not high enough to cause bursting, the membrane develops one or more inner and smaller sub-vesicle(s) engulfing the solute composition of the environment (Supplementary Movie 2b).
3. *Discocyte*: as the solute concentration *ρ*_*out*_ is decreased or the ratio *ν* is increased the vesicle instead reaches a flat plate-like shape (Supplementary Movie 2c).
4. *Elongated*: as the concentration ratio *ν* is further increased (*ν* ∼ 0.6), the vesicle transitions to a prolate configuration, closer to an ellipsoid (Supplementary Movie 2d).
5. *Globular*: for moderate values of the concentration ratio (0.6 ;S *ν* ;S 0.75) the vesicle is distorted from a normal spherical shape but doesn’t present a clear symmetry break (Supplementary Movie 2e).
6. *Sphere*: at high concentration ratio (*ν* ∼ 1) the vesicle conserves its sphere-like fluctuating shape independently of the solute concentration *ρ*_*out*_ (Supplementary Movie 2f).

A few important conclusions can be taken from Figure 3: 1) We recover the main equilibrium shapes – namely prolates (labeled here as globular or elongated), discocytes and stomatocytes – observed experimentally [9, 15], numerically [32, 33], and predicted theoretically [26–28] 2) Importantly, we identify a large subset of the parameter space where the osmotic shock leads the vesicle to inner compartmentalization (labeled as *Inner bud* in the figures), absorbing exclusively external solute particle into a smaller sub-vesicle inside the original one. Such a process is very similar to experimentally-observed endocytic events that could be relevant to how proto-cells first developed compartmentalization as a form of reaction tanks or proto-organelles in the early stages of life evolution [14, 15]. 3) The dependence of the final morphology proves to be highly non-trivial and does not only depend on the concentrations ratio *ν*, which would lead to a diagram consisting only of horizontal lines as equilibrium curvature models predict [26, 27] (see Section V in the Supplementary Information). We thus find that assuming an equilibration of the osmotic shock only via volume adaptation, as previously considered, is not sufficient to properly describe shape transformations in vesicles. In order to unveil the underlying process, we now turn to the dynamics of the system and study how the morphological changes of the vesicle (characterized by its shape, surface area and volume) dynamically relate to the relaxation of the osmotic pressure across the membrane.

### C. Surface stretching and volume adaptation are coupled to the shock magnitude

Consistent with previous work [9, 15, 27, 28], we find that, upon osmotic shock, the vesicle deforms by adapting its volume to reduce the pressure on its surface Π. The previous theoretical work assuming constant surface area and a perfect osmotic term cancellation (see Section V in the Supplementary Information^†^) predicts that the equilibrium shape and its reduced volume, v_*th*_, will depend solely on the ratio of the solute concentrations on the inner and outer side of the membrane, such that v_*th*_ = *ρ*_*in*_*/ρ*_*out*_. Here the reduced area is defined as the ratio of the equilibrated and initial surface areas, a_*τ*_ = *A*_*τ*_ */A*_0_, and analogously the reduced volume as v_*τ*_ = *V*_*τ*_ */V*_0_ (see Sections I and II in the Supplementary Information^†^ for more details on the computation of these quantities). However, as shown in Figure 4a, we find that the relaxed equilibrium volume does not follow exactly the concentration ratio imposed by the osmotic shock, like theoretical considerations based purely on membrane bending energy and constant surface area would predict [27, 28]. The reason behind this discrepancy is the fact that a part of the osmotic shock in our model is absorbed by membrane stretching. This is visible in Figure 4b, where it is apparent that a change in volume is associated with the change in the surface area. Note that, because solute particles freely diffuse in the system, the dynamics is set by the diffusion timescale (*τ*_*D*_ = 100 time steps here), which corresponds to the average time needed for diffusive particles to travel one unit length on average, and sets the lower limit for the right regime of the system. This means that membrane and solute particles do not interact before this time and explains why no change is observed in Figure 4b over the initial 100 time steps.

**FIG. 4:**
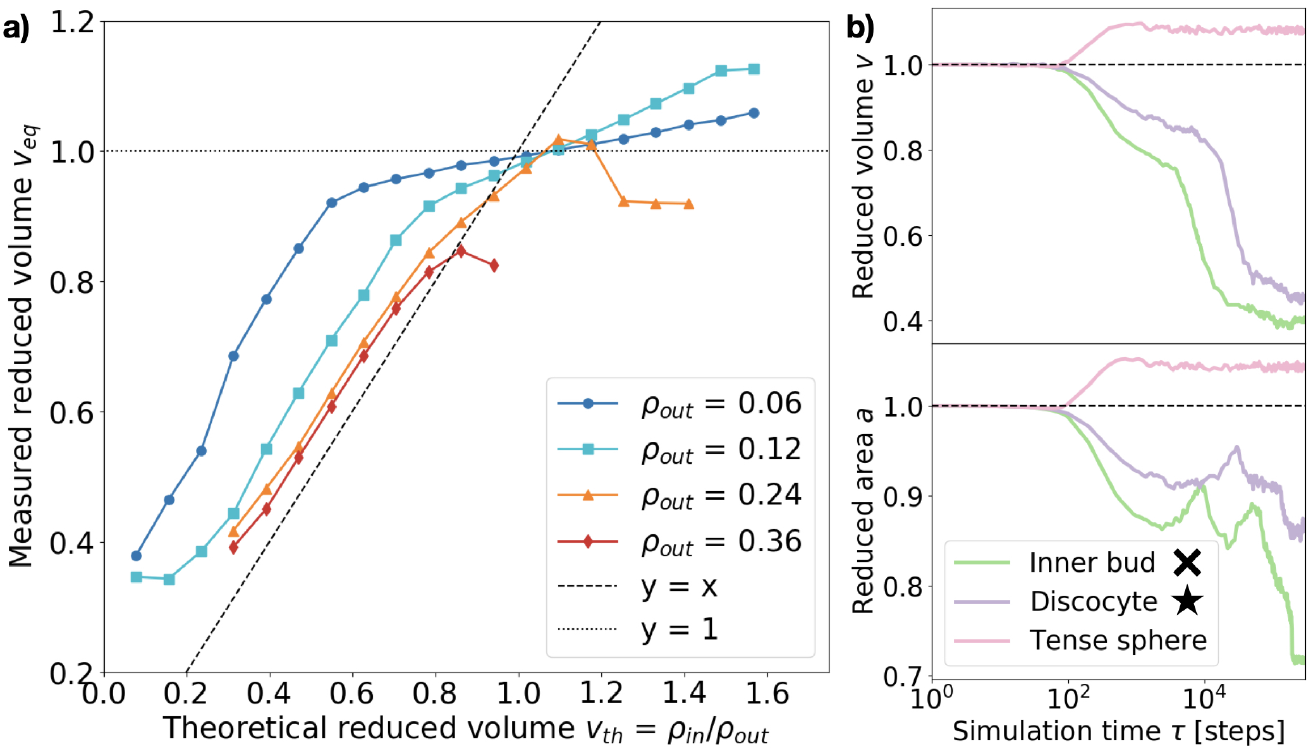
Equilibrium reduced volume and relaxation dynamics of the volume and surface area. (a) Resulting reduced volume v_*eq*_ after relaxation plotted against the expected value v_*th*_ = *ρ*_*in*_*/ρ*_*out*_. Data points are colored according to the outer particle density *ρ*_*out*_ (see legend). The dashed line represents *y* = *x* and the dotted line *y* = 1. Each point corresponds to the equilibrium average over 5 independent runs. Error bars are negligible. Reduced surface area a and reduced volume v relaxation curves for three representative cases (see legend).<u>Inner bud</u>: *ρ*_*out*_ = 0.18 *ν* = 0.31. <u>Discocyte</u>: *ρ*_*out*_ = 0.12 *ν* = 0.31. <u>Tense sphere</u>: *ρ*_*out*_ = 0.18 *ν* = 1.25. The simulation time *τ* on the x-axis is in logarithmic scale and expressed in time steps. The cross and star symbols refer to the corresponding points on the morphology diagram (see Figure 3).

As Figure 4a shows, the actual reduced volume does scale linearly with the theoretically expected one for low values (v_*th*_ *<* 1) but instead remains always above it and saturates around v_*th*_ = 1. For v_*th*_ *<* 1, the difference between the expected and observed values overall increases as the solute concentration decreases; for v_*th*_ *>* 1, instead, v_*eq*_ does not exactly collapse to one and actually exceeds it. This observation, together with the membrane area change displayed in Fig.4b, strongly suggests that a part of the energy provided by the osmotic shock is transformed into area strain, coupling it to the volume evolution in a non-trivial way. This indicates that volume and area relaxation is an intricate process where the equilibrium volume does not only depend on the ratio of inner and outer concentrations (v_*eq*_ ≠ *v*_*th*_ = *ρ*_*in*_*/ρ*_*out*_) but rather on the actual magnitude of the osmotic shock.

Membrane stretching is rarely accounted for in theoretical models, which does not allow such models to explore hypotonic conditions. Molecular dynamics simulations can access these conditions as shown by the magenta curves in Figures 4b and 5, corresponding to a *Tense sphere*: the vesicle maintains its spherical shape but increases its volume via stretching of the membrane. Nevertheless, when assessing the simulation results it is important to consider how the mechanical properties of the simulated membrane compare to those of biologically relevant membranes. In the case simulated here, the compressibility modulus of the membrane is estimated to be around 1 mN/m (assuming the membrane bead size *σ* ∼ 10 nm) [34] which is around an order of magnitude lower than, for instance, the three-beads-per-lipid CG lipid membrane model [36] and even further from experimentally measured values for single bilayer vesicles of varying composition (ranging from ∼ 60 to ∼ 1700 mN/m) [37, 38]. However, the rupture tension (tension beyond which the membrane breaks) presents similar disparities, resulting in critical area strains a_*c*_ (normalized surface area change beyond which the membrane breaks) that are consistent with the experimental ones. Indeed, for the model used in this work a_*c*_ 0.09 [34] (very similar to that of the Cooke *et al*. membrane model [36] of a_*c*_ ∼ 0.08) while experiments have measured typical lipid bilayer vesicles to have a critical area strain of a_*c*_ ∼ 0.02 − 0.05 [37, 38]. This suggests that our simulations are somewhat overestimating membrane stretching (9% instead of 5%) but does not invalidate the role of membrane stretching in the response to osmotic shock. In fact, membrane stretching in response to osmotic shock is known to be crucial for the equilibration of osmotic pressures at the cell interface via mechanosensitive channels in unicellular organisms [39, 40].

Taken together, the above observations indicate that the absorption of osmotic shocks by volume adaptation is more involved than just a volume equilibration at a constant surface area. Indeed, we find that membrane stretching plays a non-negligible role and absorbs part of the energetic shock. Moreover, the magnitude of the volume change depends on the magnitude of the shock, which in turn is dependent on the solute concentration: for a same concentration ratio, the larger the absolute solute concentration the larger the deformation.

### D. Osmotic shock adaptation is a two-step process: absorption and dissipation

Next we turn to the dynamics of the relaxation process. We observe that the system displays a striking separation of scales in the relaxation process towards equilibrium, as visible in Figure 5a. This Figure shows the measured osmotic pressure and vesicle volume (see Section I in the Supplementary Information^†^) for three representative cases: i) a hypertonic osmotic shock resulting in an inner sub-vesicle (green line, indicated with a cross on the diagram in Figure 3), ii) a hypertonic shock resulting in a discocyte (purple line, indicated with a star on the diagram in Figure 3) and ii) a hypotonic shock resulting in a tense sphere (pink line).

**FIG. 5:**
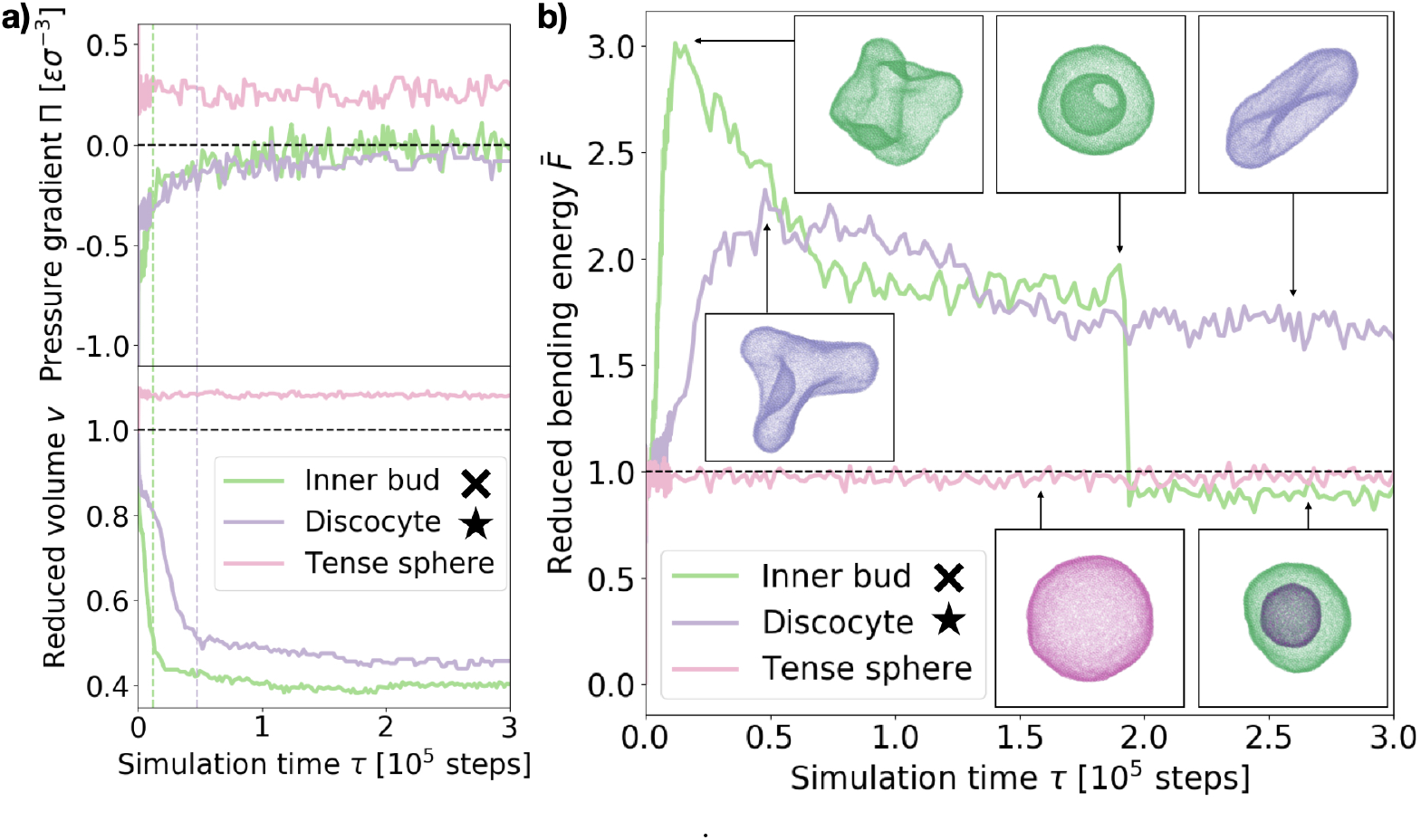
Shock relaxation dynamics. (a) Osmotic pressure difference across the membrane Π (in MD units: [Π] = *ϵσ*^−3^) and reduced volume v relaxation curves for three representative cases (see legend). <u>Inner bud</u>: *ρ*_*out*_ = 0.18 *ν* = 0.31. <u>Discocyte</u>: *ρ*_*out*_ = 0.12 *ν* = 0.31. <u>Tense sphere</u>: *ρ*_*out*_ = 0.18 *ν* = 1.25. The green and purple dashed vertical lines indicate the time of maximum bending energy for the “Inner bud” and “Discocyte” scenarios respectively (see panel (b)). (b) Bending energy evolution for the same three representative cases (see legend). The dashed line is the mean bending free energy of a relaxed spherical vesicle of the same size. *Inset*: Snapshots of the vesicles at different points in time for each scenario. Particles are colored accordingly. Time is in simulation time steps. The cross and star symbols refer to the corresponding points on the morphology diagram (see Figure 3).

Taken together, these results show the volume and pressure dynamics are coupled and occur on a much shorter time scale than the shape relaxation, since equilibrium for both Π and v is reached at times shorter than *τ* ≪10^5^ time steps in all cases. Interestingly, such volume dynamics display striking similarities with the osmotic shock relaxation curves presented in the work by Gabba *et al*., where the authors have observed an analogous decaying profile in the normalized volume evolution of impermeable vesicles subject to hypertonic osmotic shocks, both *in vitro* [29] and in a physiochemical kinetic model [30]. The hypotonic scenario resulting in a tense sphere is of particular interest, as it presents an incomplete osmotic pressure relaxation: Π_*eq*_ ∼ 0.25 *σ*^−3^. This is caused by the vesicle reaching its maximal allowed volume (for maximal stretching of the lipid membrane, a_*c*_ 0.09) before the pressure difference is completely cancelled. However, this remaining osmotic stress is not sufficient to overcome the energy barrier for bursting the vesicle, allowing for the observed morphology. For hypotonic shocks of larger magnitude we observe vesicle bursting instead of the tense sphere morphology. This is a distinct feature of hypotonic scenarios, indicating that vesicles are inherently much better prepared to endure hypertonic shocks. A possible way of adaptation to such conditions is the successive opening and closing of pores to release the tension imposed by the solution, which has been observed experimentally [24]. However, our simulations do not recover such cyclic dynamics, for which we believe a more detailed membrane model would be necessary.

Tracing the behaviour of the mechanical energy of the vesicle during the relaxation process, measured by the reduced bending free energy 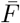 in time (refer to Section III in the Supporting Information for a detailed definition and computation details), allows to gain further insight into the relaxation process (Figure 5b). For the discocyte and inner bud scenarios (purple and green curves, respectively), we observe a sharp maximum in the bending energy which is concurrent with the volume equilibration (*τ* ∼1.2 ×10^4^ time steps and *τ* ∼ 4.8 ×10^4^ time steps respectively, as shown by the two vertical lines on Figure 5a). This indicates that the osmotic shock absorption induces very strong deformations on the vesicle which are then smoothed out over time as the vesicle relaxes to its minimum energy. Notice that for the green curve in Figure 5b this implies an inward budding event which results in a sudden drop of the energy. For the tense sphere the vesicle does not change its shape and remains spherical throughout the simulation, hence the bending energy fluctuates around 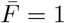. Refer to Supplementary Movies 2b, 2c and 2g for a better visualization of the relaxation process.

In the case of the vesicle that relaxes to the discocyte conformation, the equilibrium value of 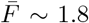 agrees very well with the value previously predicted by the analytical models that consider only vesicle curvature [27, 28]. However for the conditions when the inner bud is formed we observe a three stage evolution: first a large amount of energy is absorbed by developing a very strong deformation (peak of 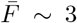), which then relaxes as 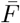 equilibrates around 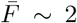 by developing a large invagination that is still connected to the parent membrane (free energy value predicted by curvature models for a stomatocyte conformation [27, 28]). Contrary to what curvature models can predict, the vesicle does not remain in this local minimum and instead evolves further towards 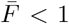 by developing an inner compartment as the invagination buds off. This *inner bud* scenario agrees very well with experimentally observed vesicle budding under osmotic stress [9, 15]. Such transformations are however inaccessible to equilibrium curvature models where vesicles are not allowed to break.

The separation of scales described in this section can thus be understood as an initial volume adaptation which absorbs the osmotic shock and a subsequent shape transformation which dissipates the accumulated tension. The first step therefore corresponds to solvent flux across the membrane while the second is achieved by lipid rearrangement. In our simulations the solvent is modelled implicitly (solute particles follow Brownian dynamics), hence the first relaxation is diffusion limited while the second one is controlled by the membrane physical properties (i.e. bending rigidity, fluidity, spontaneous curvature, compressibility modulus).

## IV. CONCLUSIONS

In this work we have adopted a novel approach in the study of vesicle osmotic shock adaptation by combining molecular dynamics with a coarse-grained lipid membrane model [34] and explicit solute particles. This allows to overcome some of the limitations of previous works while still recovering the main known features of this physical process.

In particular, here we demonstrate that with this approach we have good control over the magnitude of the osmotic shock vesicles are subject to, which scales appropriately with solute concentrations inside and outside the lipid body (see Figure 2). Moreover, we show that this method allows to investigate known equilibrium vesicle morphologies and how they relate to the osmotic perturbation. Indeed, not only do we recover the main vesicle morphologies predicted [27, 32] and observed experimentally [9] in previous works but we can also precisely map them to the relevant parameters of the system (see Figure 3).

Furthermore, using MD simulations allows for membrane scission, bursting and budding events, typically inaccessible to other modeling approaches. This enables us to investigate biologically relevant topologies such as the inner bud (or vesicle-inside-vesicle), where the perturbed vesicle develops an inner compartment engulfing external solute to accommodate the osmotic shock. In this work we thus define a subset of the parameter space for which endocytosis happens as a result of osmotic shocks (see Figure 3), possibly an important element of proto-cell development in the early stages of life [9, 14, 15]. Likewise, working with a coarse grained MD model allows to explore hypotonic shocks as well and how these result in tense spheres, with increased surface area.

Another important feature of the approach presented in this work is that it allows to track the dynamics of the relaxation process. While molecular dynamics have already been considered in previous studies of vesicular osmotic responses [33], the present work offers important advantages with respect to these in that the relaxation of the osmotic shock here is purely the result of the molecular interactions while in previous works it was imposed. We thus avoid any assumptions regarding the relaxation dynamics and work with membrane transformations defined only by the imposed osmotic shock. This allows us to dynamically characterize the transformation process by following the evolution of relevant physical magnitudes such as the bending free energy, reduced volume or pressure differential across the lipid bilayer (see Figure 5). As a result, we find that the vesicle evolution upon osmotic upshift is a two-step process where the membrane first absorbs the shock by deforming away form its spherical shape (volume and surface area changes), which results in a sharp increase of the bending energy, then dissipated by reaching the observed equilibrium morphology. Overall, the dynamic insight thus obtained provides valuable information complementing the shortcomings of classical approaches to the problem.

In the present work we have restricted our results to model membranes displaying physiologically relevant physical properties in simple scenarios. However, the combination of a coarse-grained membrane model with explicit solute particles, which allows to precisely control different aspects of the membrane physics, could easily be applied to systems of growing complexity (e.g. membranes with non-zero spontaneous curvature or in different fluid phases). Possible further applications of the approach taken here could thus be to study multiple endocytic events in fluctuating and heterogeneous environments, which could give rise to complex internal vesicle structures of different compositions. The model is also well suited for studying vesicle bursting cycles observed in hypotonic environments [24] or the different morphologies observed for multiphasic [8] and protein charged vesicles [11, 19], where the dynamical nature of MD simulations should prove useful.

## Supporting information

Supplementary Movie 2a (burst)

Supplementary Movie 2b (inner bud)

Supplementary Movie 2c (discocyte)

Supplementary Movie 2d (elongated)

Supplementary Movie 2e (globular)

Supplementary Movie 2f (relaxed sphere)

Supplementary Movie 2g (tense sphere)

## Acknowledgements

We acknowledge support from the Royal Society (C.V.C and A.Š.), the Medical Research Council (C.V.C and A.Š.), and the European Research Council (Starting grant EP/R011818/1 to A.Š.). We thank Johannes Krausser and Ivan Palaia from the Šarić lab for useful discussions.

