## Supplementary Information for "Modelling the dynamics of vesicle reshaping and scission under osmotic shocks"

This Supplementary Information provides additional details on the computational implementation of different physical measurements of the system as well as a discussion of the analytical framework within which this work is contextualized. It also includes a section devoted to the technical details of the model used for the lipid bilayers.

### I. MEASURING VOLUMES, CONCENTRATIONS AND OSMOTIC PRESSURES

Given the nature of this study, we are particularly interested in tracking the time evolution of the volume, particle concentration and pressure difference across the vesicle. However, the strong deformations vesicles are subject to in some scenarios make measuring such physical quantities highly non trivial. To sort this issue, we opt for discretizing the space and filling each volume region (i.e. each vesicle or sub-vesicle) starting from its center of mass until we reach the limit surface (the membrane enclosing it). We then integrate the physical quantities of interest (such as the volume, the particle concentration or the pressure) over the appropriately labeled cubic bins (see Equation S1).

$$
\begin{aligned}
V_i &= \sum_{j \in i} \sigma^3 & \rho_i &= \sum_{j \in i} n_j / V_i \\
P_i &= \sum_{j \in i} p_j / (\rho_i \; V_i) & \Pi_i &= P_i - P_{i+1}
\end{aligned} \tag{S1}
$$

In Equation S1, $i$ indicates the label of the region and $j$ runs over cubic bins. $n_j \in [0,1]$ is the number of particles in bin $j$ while $p_j$ is the local pressure on that particle. The procedure is the following:

1. **Discretizing the space**: The simulation box is homogeneously divided into cubic bins of size $l = \sigma$ and volume $V_j = \sigma^3$. This choice of bin size gives the maximum precision possible.
2. **Counting particles**: This discretization allows to build a 3D matrix of occupancy numbers $n_j \in [0,1]$ according to the particle positions. For example, if a solute particle is at position $[0.5, 0.1, 0.2]$ then $n_j = 1$ for bin $j = 1$ with sides $x, y, z \in [0, \sigma]$.
3. **Local pressures**: A second matrix containing local pressure values is built similarly by computing the trace of the stress tensor on the corresponding particle: $p_j = 1/(6\pi\sigma^3)(S_j^{xx} + S_j^{yy} + S_j^{zz})$ where $S$ is the stress tensor. This local stress tensor includes a kinetic term as well as a virial term, taking into account the interaction forces between particles, and is provided by the software (LAMMPS [1]) for each particle as the simulation runs. For more technical details on how this is done see the LAMMPS documentation [2] or the original publication where it was introduced [3].

Having matrix representations of the physical quantities of interest, it only remains to properly define the domains corresponding to the different volumes of the system (inside and outside the vesicle) over which we need to integrate the local variables. For this purpose we start evaluating bins at the center of mass of the vesicle particles and fill the space by expanding in all directions via close-neighbor bins

*Electronic address:

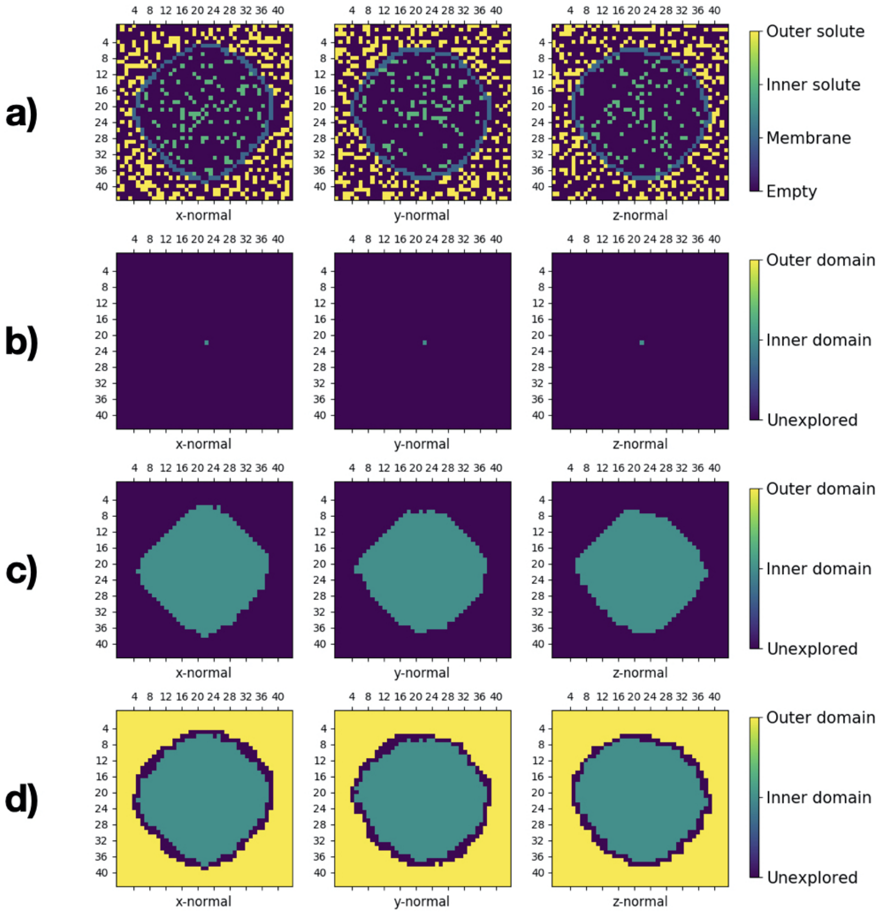


FIG. S1: **Spatial analysis of the system for $\rho_{out} = 0.3\ \sigma^{-3}$ and $\nu = 0.4$ at $\tau = 2e3$ time-steps.** *Top row* (a): Three perpendicular slices (x-, y- or z-normal) of the 3D matrix with bins color coded according to particle type. *Bottom three rows* (b-d): Three perpendicular slices (x-, y- or z-normal) of the 3D matrix with bins color coded according to volume classification. Row (b) corresponds to the initial exploration step, considering only the center of mass bin. Row (c) corresponds to a later exploration step where some parts of the interface have been reached already but the inner volume hasn't been fully explored yet. Row (d) corresponds to the final exploration step where both volumes have been explored completely and all bins in the system are appropriately classified.

until reaching a bin containing membrane particles (see rows b) and c) in Fig. S1). However, the membrane particles are not always homogeneously distributed over the surface they define, with some areas occasionally being over-stretched. This leads to artificial holes in the binned surface which we need to take into account. We do so by evaluating the set of 3D closest neighbours and neglecting individual one-bin-sized holes that are completely surrounded by a fully connected set of membrane bins. After successively implementing this protocol we reach a point where all neighbors of the explored volume are membrane bins. We then change domains and continue exploring starting from these. Applying this procedure repeatedly allows to easily classify the three-dimensional matrix bins according to their volume domain independently of membrane morphology (see row d) in Fig. S1 or Supplementary Movie 1 for a complete animation of this process).

### II. MEASURING SURFACE AREAS

Besides volumes and other three-dimensional domain physical quantities we are also interested in characterizing the stretching of the vesicle. To do so we track the temporal evolution of its surface area throughout the simulation via a particle based surface construction algorithm [4] available with the Ovito visualization software [5] (see Fig. S2). The basic principle behind this method is to generate an

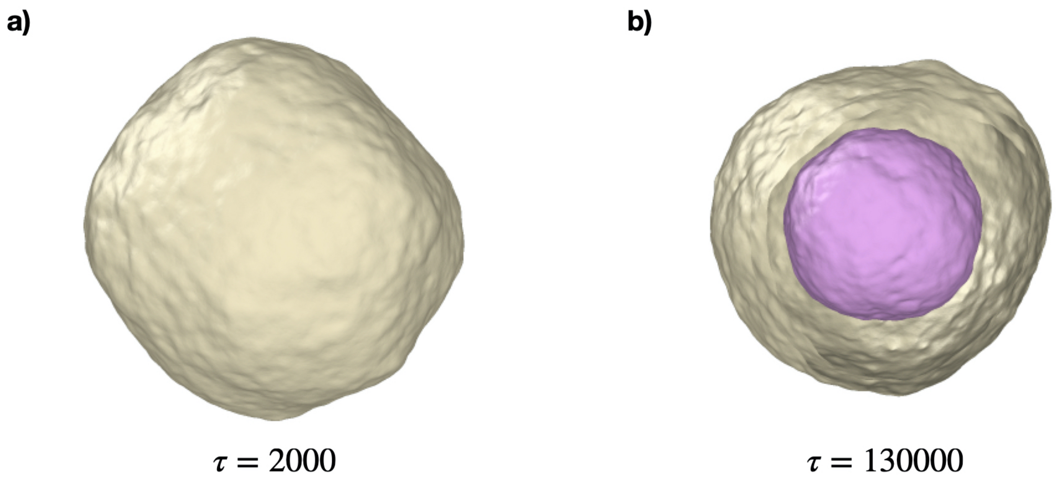


FIG. S2: Surfaces generated via the gaussian-density algorithm [4] for vesicles at different time-steps in a system with $\rho_{out} = 0.3\ \sigma^{-3}$ and $\nu = 0.4$. Note that the algorithm generates two isosurfaces per vesicle (one on the inside and one on the outside of the lipid bilayer). We thus estimate the real surface area to be the mean between the two $A_{vesicle} = \frac{1}{2}(A_{in} + A_{out})$. (a): $\tau = 2e3$ time-steps. (b): $\tau = 1.3e5$ time-steps

isosurface of the density field computed from the superposition of 3-D Gaussian functions placed at each particle site [4]. Note that this method generates two isosurfaces per cluster of membrane particles (one on the inside and one on the outside, see right panel in Fig. S2), so we take the mean of these two values as the estimate of the real surface area of the corresponding vesicle.

### III. MEASURING CURVATURES AND BENDING ENERGY

A highly interesting physical property of the vesicle for these systems is the local curvature of the membrane and its related free energy, especially in the context of the curvature models developed since Helfrich [6]. In this work we thus characterize the mechanical energy of a vesicle via its reduced bending free energy $\bar{F}$.

Given that the bending free energy $F$ of a membrane body with vanishing spontaneous curvature (like the kind we are studying here) is defined as $F = \int (2\ \kappa\ H^2 + \kappa_G\ K^G)\ dA$ (where $H$ is the local mean curvature, $K^G$ the local Gaussian curvature and $\kappa$ and $\kappa_G$ the bending rigidities of the membrane) and generally $\kappa = -\kappa_G$ for biologically relevant membranes [7–10], in this work we evaluate the bending free energy from measuring the average of the local mean curvature $\langle H_i^2 \rangle_i$ minus the ensemble average of the local Gaussian curvature $\langle K_i^G \rangle_i$, normalized by the same quantity for a relaxed free vesicle without solutes.

From the simulation we can easily determine the two principal local curvatures $C_1^i$ and $C_2^i$ at the position of a particle $i$ by computing the gradient of the normal vector to the surface along the $N_{neighs} = 6$ closest neighbours (see section below for a detailed discussion on the method used here). Having the two principal curvatures, computing the local mean and Gaussian curvatures is fairly straightforward: $H_i = 0.5\ (C_1^i + C_2^i)$ and $K_i^G = C_1^i\ C_2^i$. We thus define $\bar{F} = (\langle H_i^2 \rangle_i - \langle K_i^G \rangle_i)/F_0$, where $F_0 = \langle H_i^2 \rangle_i - \langle K_i^G \rangle_i$ for a relaxed vesicle of the same size at equilibrium (i.e. not subject to an osmotic shock). Working with this reduced bending free energy $\bar{F}$ allows to compare easily with theoretical predictions for $F_b/(8\pi\kappa)$ such as in Seifert *et al.* [11] without worrying about converting from MD to SI units.

### IV. NORMAL TO THE MEMBRANE SURFACE COMPUTATION

In the present membrane model [12] the dipole moment of membrane particles is subject to fast rotational dynamics, which leads to some fluctuations around the local normal to the membrane surface. This could lead to considerable errors when computing local curvatures from the gradient of normal vectors (see the above section). To address this issue we smooth the fluctuations out by taking the average over a set of particles instead of considering just a single dipole. Given that the membrane particles tend to organize on a hexagonal lattice, when computing the normal at the position of particle $i$ we average over its $N_{neighs} = 6$ closest neighbors. However, we found that just implementing a simple arithmetic average resulted in the introduction of artificially parallel normal vectors. In order to avoid this phenomenon we decided to work instead with Gaussian weighted averages where each neighbor's dipole is weighted according to its distance to the particle of interest (see Equation S2, where $\vec{n}_i$ is the normal vector to the surface at the position of particle $i$, $\vec{\mu}_i$ is the dipole moment of particle $i$ and $\vec{r}_i$ its position.). After trying different averaging protocols we found that this was the method that gave the lowest errors in curvature values for known membrane geometries (e.g. tubular or spherical).

$$\vec{n}_i = \frac{\vec{\mu}_i + \sum_{j=1}^{6} w_j \ \vec{\mu}_j}{|\vec{\mu}_i + \sum_{j=1}^{6} w_j \ \vec{\mu}_j|} \qquad\qquad w_j = exp(-|\vec{r_i} - \vec{r_j}|^2) \tag{S2}$$

### V. CURVATURE MODELS

Starting with Helfrich in 1973 [6] there have been a great number of developments concerning analytical models for lipid membrane deformations. The so-called curvature models thus emerged, describing vesicle deformations by minimizing free energies defined around the curvature changes of the membrane [11, 13]. In this section we will discuss how such models rely on overly simplistic approximations for the volume and area relaxation in response to osmotic shocks.

In his seminal paper [6], Helfrich defined the curvature or bending free energy per unit area as $f_b = 2\ \kappa\ (H - C_0)^2 + \kappa_G\ K^G$, where $H$ and $K^G$ are the mean and Gaussian curvatures of the bilayer respectively, $C_0$ the spontaneous curvature and $\kappa$ and $\kappa_G$ are the bending rigidities of the membrane. This is the first term in the free energy computation of a vesicle conformation, which corresponds to the cost of deforming the membrane away from its natural shape, given by $C_0$.

When considering an osmotic shock scenario, a second term needs to be introduced to account for the osmotic pressure the solute particles exert on the vesicle. This is given by $\Pi = RT\ (n/V - \rho_{out})$ where $R$ is the gas constant, $T$ the temperature, $n$ the number of moles of solute inside the vesicle, $V$ the volume of the vesicle and $\rho_{out}$ the solute concentration in the medium (assumed constant for sufficiently large bath volumes) [13].
Finally a third term can be introduced to account for the cost of stretching the membrane as a result of the deformations, although this last term is usually neglected in analytical calculations due to the small area strain biological membranes can generally admit. As introduced by Helfrich in his original paper [6], such a term should be quadratic in the relative change in area $\Delta A/A_0$: $f_s = \frac{K_A}{2}\ (\Delta A/A_0)^2$.

$$F = \int_{A_0}^{A_{eq}} f_c\ dA + \int_{V_0}^{V_{eq}} \Pi\ dV + \int_{A_0}^{A_{eq}} f_s\ dA \tag{S3}$$

Consequently, the complete free energy can be expressed as presented in Equation S3, where the integration limits are given by the initial or relaxed conformation and the equilibrium configuration respectively. An important consideration arises, since, as discussed by Seifert previously [13], there are two distinct energy scales in this equation: the first term scales with the bending rigidities ($\kappa$ and $\kappa_G$) while the second and third terms scale with the vesicle's surface area and volume respectively thus being several orders of magnitude over the curvature term. This implies that in order to solve the shape equations for the equilibrium configuration one must solve the two scales independently.

As a result, analytical models of this kind generally assume constant surface area ($A_{eq} = A_0$), which cancels the stretching term in Equation S3, and perfect volume equilibration ($V_{eq} = n/\rho_{out}$) thus canceling the osmotic pressure term as well. Such constraints can be expressed in terms of the reduced area and volume (better related to the results presented in the main text) as $\mathrm{a}_{eq} = A_{eq}/A_0 = 1$ and

$v_{eq} = V_{eq}/V_0 = \rho_{in}/\rho_{out}$, where we define $\rho_{in} = n/V_0$ the initial solute concentration inside the vesicle.

These equilibrium constraints are thus a simplification that enables achieving analytical solutions for the model but neglect the possible interplay between the area, volume and shape relaxations, providing little physical insight into the actual relaxation process as a result.

### VI. DETAILS ON THE MEMBRANE MODEL USED

In this work we have simulated lipid membrane vesicles following the single particle thick model developed by Yuan *et al.* [12]. This model is able to capture physiologically relevant properties of lipid bilayers in a relatively simple and computationally manageable representation by coarse-graining multiple lipid molecules into single beads representing sections of the bilayer around 10 nm in width. Furthermore, each of these beads is associated to a vector representing the average orientation of lipid molecules (or the normal to the membrane) at that point. The membrane beads interact with one another via a specific attractive potential that depends not only on the distance between beads but also on the relative orientation of their vectors, thus capturing curvature effects as well as fluidity, bending and other crucial aspects of membrane physics. Finally, it being a particle-based model (albeit a very coarse-grained one) it is capable of reproducing fission and fusion events, difficult to capture in simulation otherwise, and it can easily be coupled with other interactions (such as the volume exclusion of solute particles in this case). This fact, together with its simplicity and computational economy relative to other particle-based membrane models, makes the model by Yuan *et al.* an ideal choice for the systems we consider in this work.

The specific shape of the interaction potential used here, given by Equation S4, is governed by four independent parameters, each related to different properties of the bilayer:

1. $\theta_0$ corresponds to the preferred relative orientation between beads and is thus directly related to the membrane's spontaneous curvature.

2. $\mu$, being the factor in front of the orientation term in Equation S4, corresponds to the energy penalty for diverging away form the spontaneous curvature and thus is related to the bending rigidity of the membrane.

3. $\zeta$, being part of the exponent of the cosine function in Equation S4, controls the slope of the attractive branch of the potential and is therefore related to the fluidity of the membrane or diffusivity of its lipids.

4. $\epsilon$ controls the energy scale of the minimum of the potential and therefore the strength of the attraction. Consequently, this parameter is related to various physical properties of the membrane, such as its fluidity, compressibility modulus or critical area strain.

In this work we set the value of these parameters to encode for a membrane of vanishing spontaneous curvature and bending rigidity of 15 $k_BT$, $k_B$ being the Boltzmann's constant and $T$ being the temperature: $\epsilon = 4.34\ k_BT$, $\zeta = 4$, $\mu = 3$, $\theta_0 = 0$. Furthermore, following the original implementation of this model, we also fix $r_{min} = 1.12\ \sigma$ and $r_c = 2.6$ to ensure proper volume exclusion (no overlapping) between beads and sufficiently long-ranged interactions. The only other parameter involved in our modelling approach is the strength of the Weeks-Chandler-Anderson potential used for solute-solute and solute-membrane interactions, which we set to $\epsilon_{WCA} = 2\ k_BT$ to ensure proper volume exclusion.

It is important to note that the above discussion is intended to provide a reference of the basic principles behind the model developed by Yuan *et al.* to help understand how and why we have implemented it in this work. However, the relation between physical properties and model parameters is more involved than we can cover here (the membrane phase itself is a result of the interplay between $\epsilon$ and $\zeta$ rather than being controlled by a single parameter, for example) and we here refer the reader to the original publication [12] for further details.

$$U(\mathbf{r}_{ij}, \mathbf{n}_i, \mathbf{n}_j) = \begin{cases} u_R(r) + [1 - \phi(\hat{\mathbf{r}}_{ij}, \mathbf{n}_i, \mathbf{n}_j)]\epsilon, \ r < r_{min} \\ u_A(r)\phi(\hat{\mathbf{r}}_{ij}, \mathbf{n}_i, \mathbf{n}_j), \ r_{min} < r < r_c \end{cases} \tag{S4}$$

$$u_R(r) = \epsilon \left[ \left( \frac{r_{min}}{r} \right)^4 - 2 \left( \frac{r_{min}}{r} \right)^2 \right] \qquad \phi = 1 + \mu[a(\hat{\mathbf{r}}_{ij}, \mathbf{n}_i, \mathbf{n}_j) - 1]$$

$$u_A(r) = -2\epsilon \left[ \frac{\pi}{2} \frac{(r - r_{min})}{(r_c - r_{min})} \right] \qquad a = (\mathbf{n}_i \times \hat{\mathbf{r}}_{ij}) \cdot (\mathbf{n}_j \times \hat{\mathbf{r}}_{ij}) + sin\ \theta_0(\mathbf{n}_j - \mathbf{n}_i) \cdot \hat{\mathbf{r}}_{ij} - sin^2\theta_0$$

### VII. SYSTEM SIZE CONSIDERATIONS

For the bulk of this work we have considered vesicles of N = 4322 model particles, each of $\sigma \sim 10$ nm in diameter, meaning that vesicles in iso-osmotic conditions have a diameter of a few hundred nanometers ($r_{vesicle} \sim 17\sigma$). While this is within the range of typical sizes for lipid vesicles, GUVs (Giant Unilamellar Vesicles) generally have diameters closer to the micron. However, simulating systems of such sizes for the full analysis presented here would have been impossible from a computational standpoint. Instead, in this section we present a few representative simulations of larger systems in order to check that are results are not affected by the system size on the scales considered.

In particular, for the larger simulations we consider vesicles made of N = 47921 model particles, resulting in an equilibrium diameter of around 1.1 $\mu$m, and apply an osmotic shock (see Section 2 in the main text) for relevant points of the parameter space and compare these results with the ones obtained for the smaller system. We thus recover the main morphology transformations discussed in the main text and observe the same separation of scales in the relaxation process. Indeed, when applying a similar osmotic shock to the larger vesicles these evolve towards equilibrium shapes equivalent to those recorded for the smaller system, as shown in Figure S3 where four representative morphologies are displayed together with their smaller counterpart. Note in particular how inner compartmentalisation events also occur for strong hypertonic shocks while hypotonic shocks result in dilated or even burst vesicles.

Furthermore, the larger system simulations display a separation of scales in the dynamics strikingly similar to that discussed in the main text for the smaller system. Not surprisingly, we observe here that volume relaxation occurs very rapidly (equilibrium is reached in the first few thousand time steps) while the smoothing of the deformations resulting from it occurs on a much longer scale ($\geq 10^6$ time steps). Hence, although the complete system relaxation takes longer for the larger system, the distinctive two-step dynamics of shock absorption and dissipation observed for the smaller system (see Section 3.4 in the main text) is still present at this scale.

Together, the above findings clearly indicate that the shape transformations of our model vesicles and the dynamics they follow remain qualitatively unchanged irrespective of the system size. Consequently, we believe that the observations presented in the main text of this work arise from the general physical properties of lipid bilayers and are applicable to vesicles of any physiologically relevant size.


[1] http://lammps.sandia.gov.

[2] https://lammps.sandia.gov/doc/compute_stress_atom.html.

[3] Aidan P. Thompson, Steven J. Plimpton, and William Mattson. General formulation of pressure and stress tensor for arbitrary many-body interaction potentials under periodic boundary conditions. *The Journal of Chemical Physics*, 131(15):154107, 2009.

[4] Michael Krone, John Stone, Thomas Ertl, and Klaus Schulten. Fast Visualization of Gaussian Density Surfaces for Molecular Dynamics and Particle System Trajectories. In Miriah Meyer and Tino Weinkaufs, editors, *EuroVis - Short Papers*. The Eurographics Association, 2012.

[5] Alexander Stukowski. Visualization and analysis of atomistic simulation data with OVITO-the Open Visualization Tool. *MODELLING AND SIMULATION IN MATERIALS SCIENCE AND ENGINEERING*, 18(1), JAN 2010.

[6] W Helfrich. Elastic properties of lipid bilayers: Theory and possible experiments. *Zeitschrift für Naturforschung C*, 28(11-12), 1973.

[7] Jan Steinkühler, Roland L Knorr, Ziliang Zhao, Tripta Bhatia, Solveig M Bartelt, Seraphine Wegner, Rumiana Dimova, and Reinhard Lipowsky. Controlled division of cell-sized vesicles by low densities of membrane-bound proteins. *Nature Communications*, 11(1):1–20, 2020.

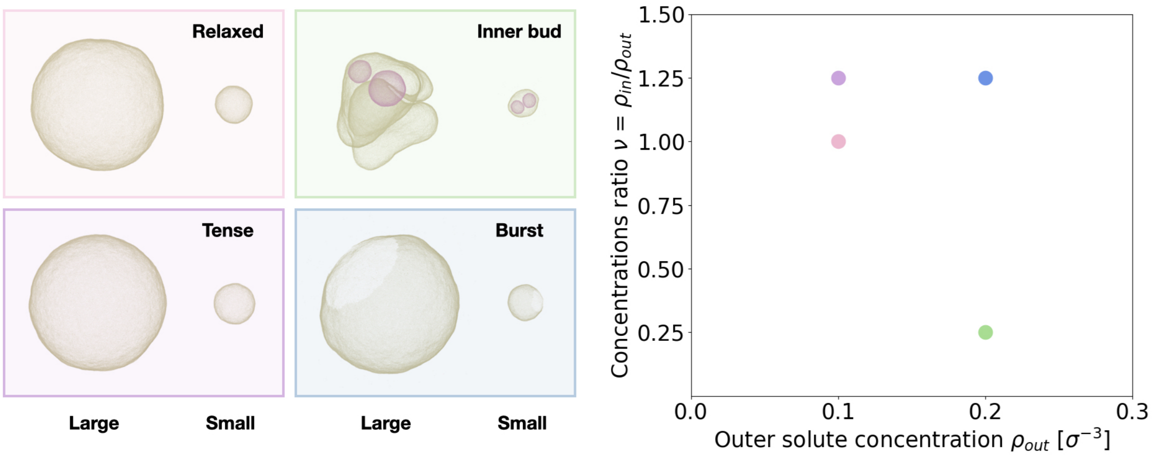


FIG. S3: Examples of equilibrium morphologies for the two system sizes considered across the parameter space. The shapes of the larger and smaller systems after equilibration of the osmotic shock are shown together (left and right of each inset respectively) for four representative scenarios (iso-, hypo- and hyperosmotic shocks), plotted on the parameter space to the right of the figure. Important morphologies such as the *Inner bud* or *Tense sphere* (see main text) are recovered.

[13] Udo Seifert. *Configurations of fluid membranes and vesicles*, volume 46. 1997.
